# Speciation across the Earth driven by global cooling in terrestrial Orchids

**DOI:** 10.1101/2021.02.06.430029

**Authors:** Jamie B. Thompson, Katie E. Davis, Harry O. Dodd, Matthew A. Wills, Nicholas K. Priest

## Abstract

Though climate change has been implicated as a major catalyst of diversification, its effects are thought to be inconsistent and much less pervasive than localised climate or the accumulation of species with time. But, we need focused analyses of highly specious clades to disentangle the consequences of climate change, geography and time. Here, we show that global cooling shapes the biodiversity of terrestrial orchids. Employing a phylogenetic framework of 1,450 species of Orchidoideae, the largest terrestrial orchid subfamily, we find that speciation rate is causally linked with historic global cooling, not time, habitation in the tropics, altitude, variation in chromosome number, or other types of historic climate change. Relative to the gradual accumulation of species with time, models specifying speciation driven by historic global cooling are 328 times more likely. Evidence ratios estimated for 212 other plant and animal groups reveal that the orchidoids represent one of the best-supported cases of temperature-spurred speciation yet reported. Employing >1.4 M georeferenced records, we find that global cooling drove contemporaneous diversification in each of the seven major orchid bioregions of the earth. With current emphasis on understanding and predicting the immediate impacts of global warming, our study provides a clear case study of the longterm impacts of global climate change on biodiversity.

**Significance statement:** The staggering biodiversity of angiosperms has been difficult to reconcile with the gradual Darwinian process thought to create it. Changes in climate through the Earth’s history could have instigated this diversification, but perceived variability across clades and geography has restrained generalisation. In this paper, we reconstruct the evolutionary history of a rich terrestrial orchid subfamily favoured by Darwin (Orchidoideae, ~5,000 species), and use >1.4 million georeferenced records to test how and where those orchid species arose. We find that global cooling between the Oligocene and present day spurred an avalanche of speciation in orchidoid assemblages across the Earth. This work resolves the orchidoid phylogeny and provides a clear example of how historic climate change drives global patterns of biodiversity.

## Main

Charles Darwin’s “abominable mystery” was why diversification can happen so rapidly (1, 2). The abrupt appearance of diverse clades of angiosperms was a challenge not only to Darwin’s theory, but to evolution itself (3). Under mounting pressure from paleobotanists and other groups, Darwin asserted an admittedly ‘wretchedly poor conjecture’ that the patterns could be explained by an ancient origin of angiosperms on a ‘small isolated continent in the southern hemisphere’ (3, letter to Joseph Hooker, 22 July 1879, Darwin Correspondence Project). Since then, there has been active inquiry into the forces determining distribution, geographic origins, and timing of diversification of all angiosperm clades (3–5).

A growing body of theory argues that the origins and contemporary distributions of biodiversity are largely determined by historic climate change (6, 7). The problem has been, however, that there are few clear supporting examples. The consequences of climate change on diversification are generally thought to be inconsistent, varying between closely related groups of taxa (8, 9), the type of climate change (10) and the ecoregion in which it occurs (11, 12). In contrast, a substantial proportion of biodiversity can be explained by gradual accumulation of species over time (13) and influences of geographic factors such as latitude (14, 15) and elevation (16, 17).

Focused analyses of cosmopolitan clades of plants are needed to disentangle the impacts of climate change from geography and time on the origins of biodiversity. Many of the classic cases of adaptive radiation, such as cichlid fish, Darwin’s finches, and *Anolis* lizards, diversification occurred at such localised scales that it is difficult to determine whether global climate change contributed to their diversification. Still, the latitudinal species gradient may not have as much of a role in speciation as previously thought (14, 15). Global warming is the most parsimonious temporal climatic explanation for diversification in many animal taxa (13). But, because animals can rapidly shift their distributions according to prevailing ecological conditions, it is difficult to exclude the impacts of localised geography. The critical unresolved question is whether historic climate change can have consistent, global influences on the origins and distributions of plant biodiversity (9, 18).

The terrestrial orchid subfamily Orchidoideae (orchidoid orchids) are an ideal clade with which to test this question. Though their recent origins (stem age c.64 MYA (19)) exclude them from contributing solutions to all aspects of Darwin’s “abominable mystery” (1–3), the orchidoids are globally distributed and extraordinarily diverse, with 5,000 species (20) and extensive distribution records. The timing of speciation events within the clade and in reference to all orders of monocots are well established, despite a scarcity of fossil evidence (19). Geographic consistency can be considered by testing for similarities in responses to climate change in taxa endemic to each of the seven major orchid bioregions (defined by Givnish et al, 21). Moreover, the clade exhibits minimal variation in traits previously associated with speciation, such as pollinia, and lacks epiphytism, which have been previously shown to contribute to accelerated speciation across all major clades of orchids (19).

## Results

### Phylogeny and Diversification

Diversification rate is varied across the Orchidoideae (Figure 1). We reconstructed a maximum likelihood phylogeny of 1,450 taxa (29% of the 5,000 known species), using nine nucleotide loci mined from Genbank. The phylogeny is well-supported, with 66% of internal nodes supported by >70% bootstrap support (BS) and 44% by >90% BS topology, which generally agrees with recent estimates (19, 22). We time-calibrated the tree with penalised likelihood (23), implementing 15 divergence constraints (Figure S1; Table S1). As there is only one orchidoid fossil, we used previously estimated secondary calibrations (19) involving plastome-level data in a Bayesian framework with an uncorrelated, relaxed lognormal clock and fossil constraints sampled across monocot orders.

**Figure 1:**
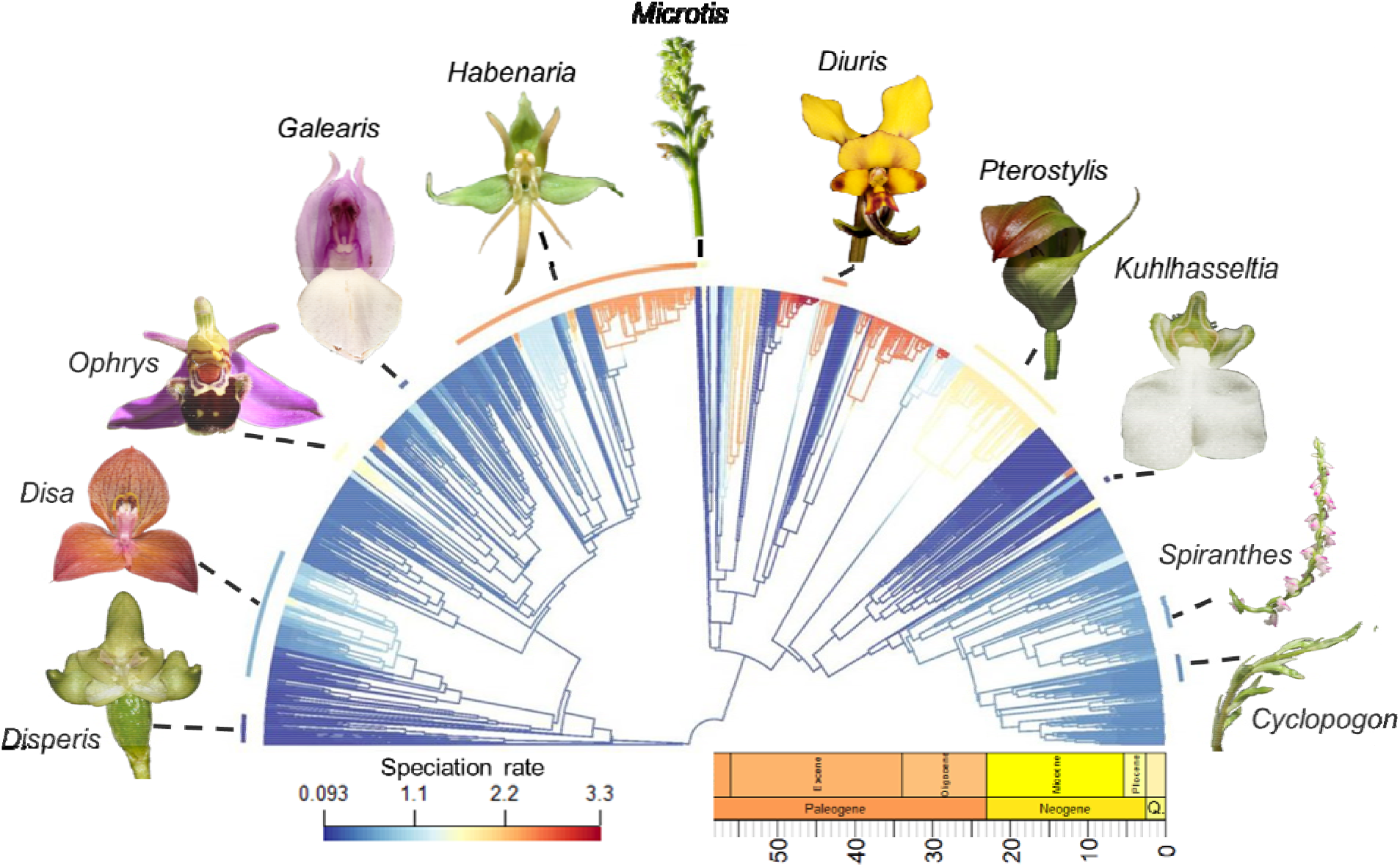
Time-calibrated phylogeny of 1,450 orchidoid taxa, visualised against the geological timescale. Branches are coloured by speciation rates estimated with Bayesian Analysis of Macroevolutionary Mixtures. Orchidoid genera representing the diversity of form are associated with arc segments coloured by their mean speciation rate. Images were sourced from Flickr (Creative Commons and modifications allowed).

We estimated speciation dynamics using the reversible-jump Markov Chain Monte Carlo (MCMC) framework Bayesian Analysis of Macroevolutionary Mixtures (BAMM) (24), accounting for incomplete sampling by specifying the known richness of each genus (20). In all analyses with BAMM, we only considered speciation rate, not extinction rate, which avoids potentially unreliable parameter estimation (25–27) and is predicted to accurately estimate speciation rate variability (28). We found 35-fold variation in the rate of speciation, with strong support for 24 rate shifts, each of which indicate significant increases in speciation rate (Bayes Factor 20.8). In the best shift configuration, the earliest rate shifts occur approximately at the Eocene-Oligocene boundary, but the majority fall between the Oligocene-Miocene boundary and the present day. These time points are relevant because they correspond with a protracted period of climatic cooling in Earth’s history (29).

### Cross-clade consistency in consequences of cooling

To formally test the hypothesis that climatic cooling drove orchidoid speciation, we correlated 9,001 realisations of the historical speciation curve with a reconstruction of Cenozoic δ^18^O (a proxy for mean global temperature) (29) and assessed whether the distribution of correlation coefficients differed from zero. Across the subfamily, speciation rate had a strong and consistently negative association with Cenozoic δ^18^O (Average of DCCA correlation coefficients = −0.50, p<0.0001; Figure 2b). Overall, we found strong evidence for a negative exponential correlation between estimated global paleoclimatic temperature data and mean BAMM-estimated speciation rates (r = −0.79, p<0.0001; Figure 3a). To control for other climatic drivers, we tested for the influence of historic values of mean atmospheric CO_2_ and global sea level variation, both of which are correlated with mean global temperature (30) and have been shown to influence speciation of other clades (ex: 10, 12, 31, 32). Though the distributions of correlation coefficients were significantly different from zero, neither atmospheric CO_2_ nor sea-level were strongly correlated with diversification in the Orchidoideae (CO2: 0.031, p<0.0001; sea-level: −0.029, p<0.0001).

**Figure 2:**
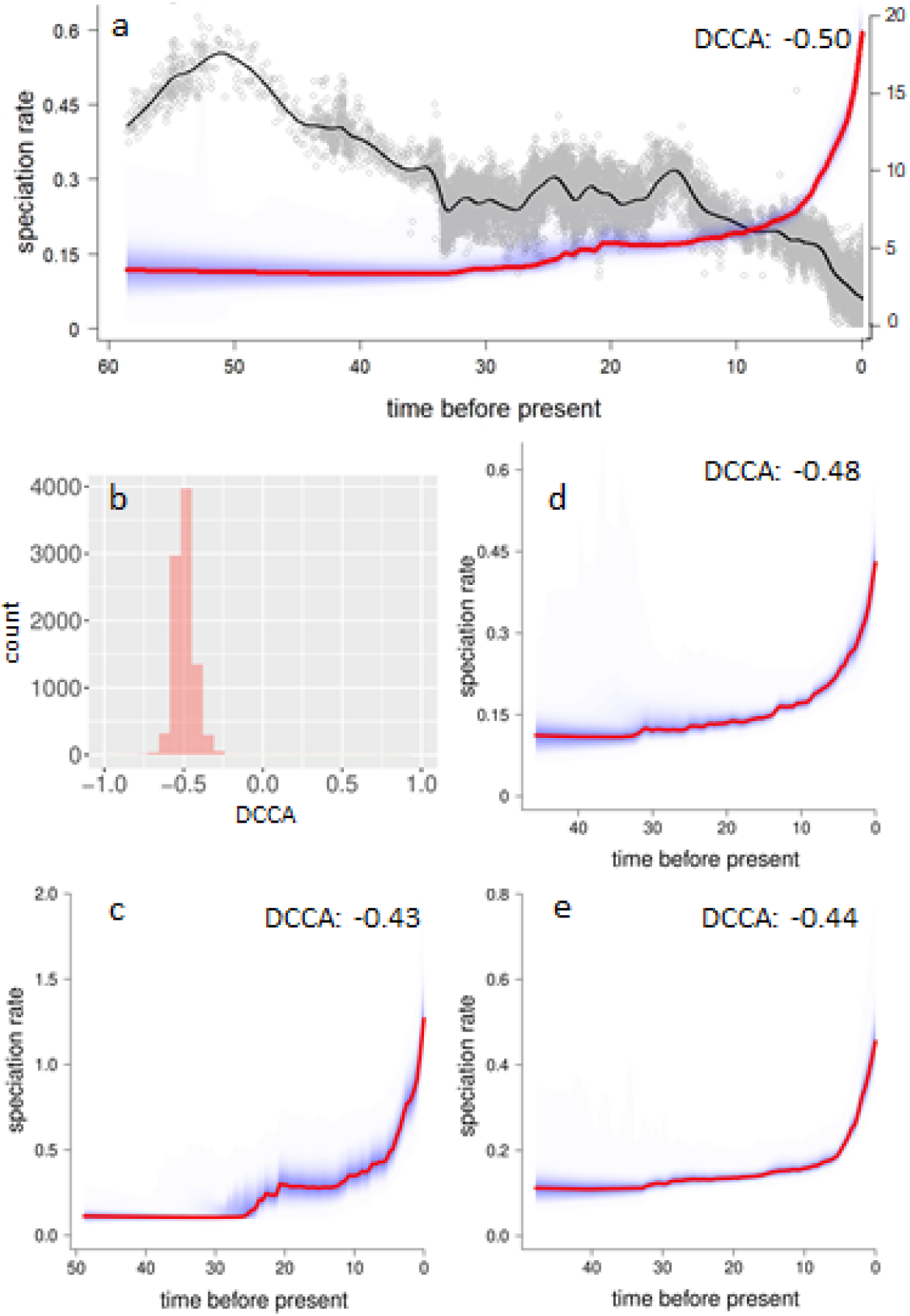
Consistency in the influence of global cooling on rapid diversification across Orchidoideae. The estimated historical speciation rate curve for the whole tree (red line with blue 95% CI) is plotted alongside the curve of estimated mean global paleotemperature (δ^18^O, black line) (**a**). A histogram of the DCCA coefficients used to infer the mean correlation coefficient across the whole tree (**b**). The estimated historical speciation rate curves of sub-clades Diurideae (**c**), Cranichideae (**d**) and Orchideae/Diseae (**e**) are reported with mean correlation coefficients indicated (all are consistently negative and significant at p<0.0001).

**Figure 3:**
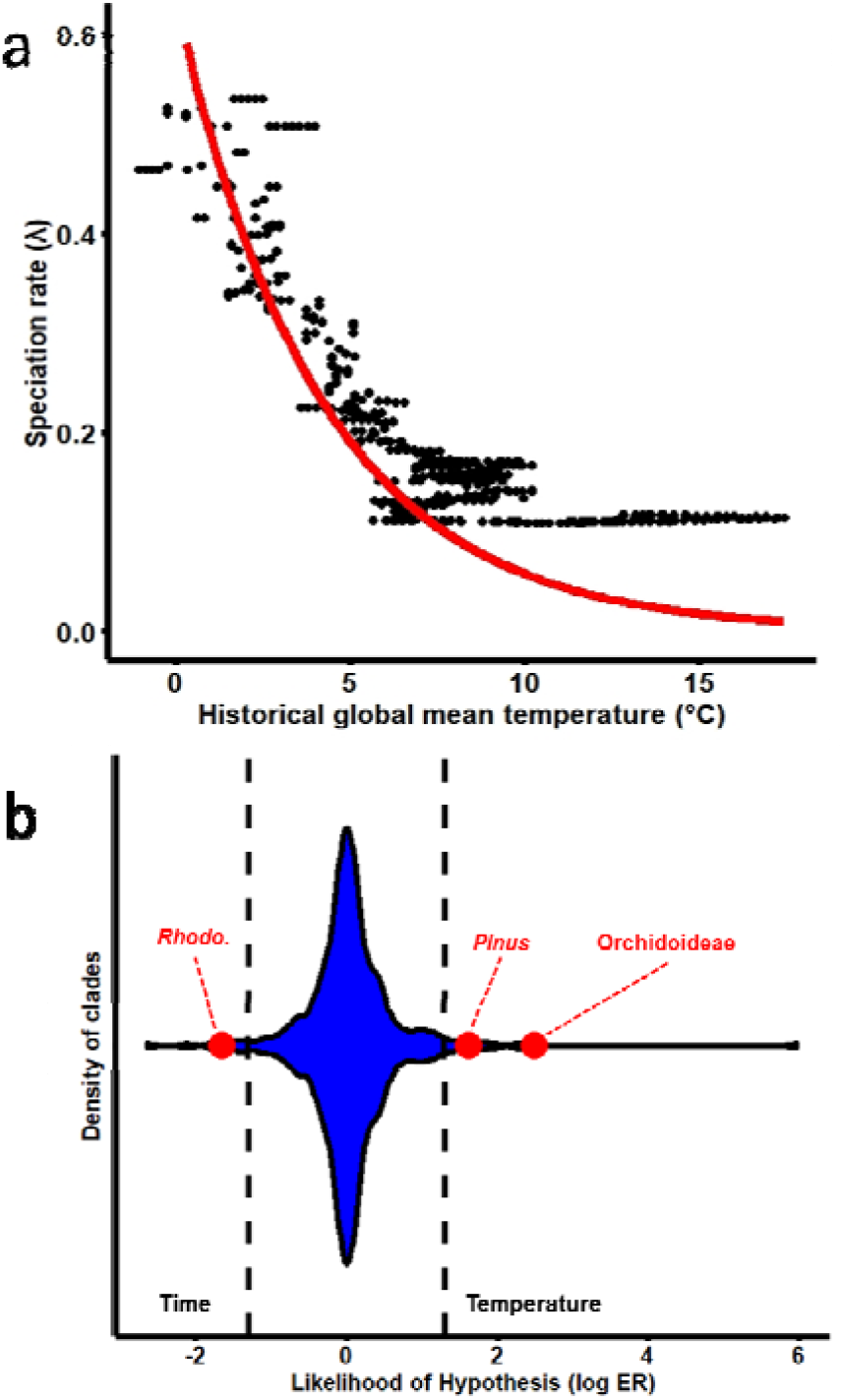
Strength of historic temperature on speciation rate. (a) The correlation between temperature and speciation rate (without estimating extinction) in the Orchidoideae is reported both for MCMC Bayesian analysis (BAMM, black dots) and likelihood-based exponential speciation model fitting (RPANDA, red curve), where speciation driven by global cooling is more likely than by time (ΔAICc = 17.68). (b) Evidential support for nested sets of temperature-dependent versus time-dependent exponential speciation models is displayed as log evidence ratios, with hatched lines indicating taxa in which time or temperature is 20 times more likely. Temperature is 328 times more likely than time to drive speciation in the Orchidoideae (red dot on right), one of the best supported examples in comparison with 210 tetrapod groups (blue violin plot) and *Rhododendron* and *Pinus* plant groups (red dots on left and in middle, respectively).

Other studies have found that the influence of climate change varies between closely related clades (8, 9). But, we found cross-clade consistency in the influence of climate change on diversification. Our DCCA analysis revealed a negative relationship between the speciation curve and historical global temperature in each of the three major subclades (Cranichideae −0.48, p<0.0001; Diurideae −0.43 p<0.0001, Orchideae/Diseae −0.44, p<0.0001; Figure 2).

Consistent responses in speciation to global cooling can’t be attributed to variation in chromosome number. With data from the Chromosome Counts Database (33), we find that each of the major orchidoid subclades have more chromosomes than earlier-diverging orchid subfamilies (Cranichideae mean = 21.52, t_256_=5.43, p<0.0001; Diurideae mean = 21.45, t_106_=4.53, p<0.0001, Orchideae/Diseae mean = 25.53, t_304_=9.31, p<0.0001). And, as a subfamily, the orchidoids (mean = 23.96 +/-0.48 SE) have more chromosomes than both earlier-diverging orchid subfamilies (Apostasioideae, Cypripedioideae & Vanilloideae mean = 15.72 +/-0.83 SE; t_227_=8.77, p<0.0001) and their more recently diverged, pollina-containing sister group (Epidendroideae mean = 22.22 +/-0.20 SE; t_628_=3.37, p=0.0008). However, after considering the similarity between chromosome counts of the sister taxa, the most parsimonious explanation is that a change in chromosome number occurred in a lineage ancestral to the epidendroids and orchidiods, well before the most recent epoch of global cooling.

Targeted analysis implicates a role for global cooling in adaptive evolution. As all orchidoids produce pollinia, packets of pollen, we were unable to test for their well-established role in accelerated speciation (19). However, diversification in the genus *Disa* is indicative of adaptive radiation because it exhibits a diversity of pollination syndromes with well-supported evidence of adaptive evolution (34, 35). Consistent with findings across the subfamily, we find that the speciation curve of *Disa* is negatively correlated with global mean temperature (DCCA = −0.46, p<0.0001).

### Temperature-, not time-, dependent speciation

For taxa exhibiting steady increases in speciation rate with time, the gradual accumulation of species with time is often found to be the most parsimonious explanation for diversification (13). Though the speciation rate of orchidoids increases up to the present (Figure 2), we find substantially stronger support for models of global cooling than time. Both MCMC Bayesian (BAMM) and likelihood-based (RPANDA (36)) model fitting approaches estimate similar relationships between historic temperature change and speciation rate (Figure 3a). In tests with RPANDA, we find that models specifying exponential influences of historic temperature on speciation fit patterns of orchids diversification substantially better than comparable time-dependent models with or without estimating exponential extinction (ΔAICc = 233.1 and 17.68, respectively). Evidence ratios for temperature- and time-specific exponential speciation (estimated for models with no, constant or exponential extinction) reveal that temperature-driven diversification is 328.3 times more probable. To assess the extent to which the orchidoids represent an archetypal example of temperature-spurred diversification, we applied this methodology to 210 animal groups (13) and two other plant groups (*Rhododendron* (37) and *Pinus* (38)). We find that the orchidoids represent one of the most convincing cases of temperature-driven speciation yet recorded (Figure 3b).

### Lack of temperature-related geographic effects

It has long been held that the tropics is a ‘cradle’ of diversity for plant and animal taxa (14, 15); but, previous simulations of global impacts of climate change on biodiversity reveal that speciation rate can appear to be geographically varied even when climate change is the central driver of that diversification (7, 39). Speciation rate appears to be geographically varied in the orchidoids. Consistent with other recent studies of angiosperm diversification (10, 14, 17, 40, 41), we find that orchidoid diversification is generally faster where it is cooler. Maps based on 1.5 million georeferenced occurrence records (42) show that the orchidoids are globally distributed with peaks of richness centred in temperate regions and a peak in speciation rate evident in temperate regions of the Southern Hemisphere (Figure 4). Though, we find no significant difference in speciation rate when defining the tropics by temperature (tropics = >18°C, non-tropics = <18°C) (SI Appendix, Fig. S1, Tab. S1), orchidoid speciation rate is significantly lower in the tropics than in temperate regions when defining the tropics by either binarised latitude (−23.5 to 23.5 degrees) or continuous latitude (SI Appendix, Fig. S2-S4, Tab. S1). However, we find no evidence of a causal relationship between geography and speciation rate, paralleling the lack of a latitudinal gradient in speciation rate seen across terrestrial orchids (19). After account for independent shifts in diversification that occur across the orchidoid phylogeny, there was no associations between latitude and speciation rate, whether by temperature (STRAPP p=0.55) (43), binarised latitude (STRAPP p=0.48), or continuous latitude (Es-Sim p=0.18, STRAPP p=0.37) (44), despite strong influences of historic global cooling on diversification. This implies that while latitudinal variation in speciation rate is apparent, the origin of new orchidiod species was primarily driven by global cooling, which cannot be explained by geographic influences on speciation rate.

**Figure 4:**
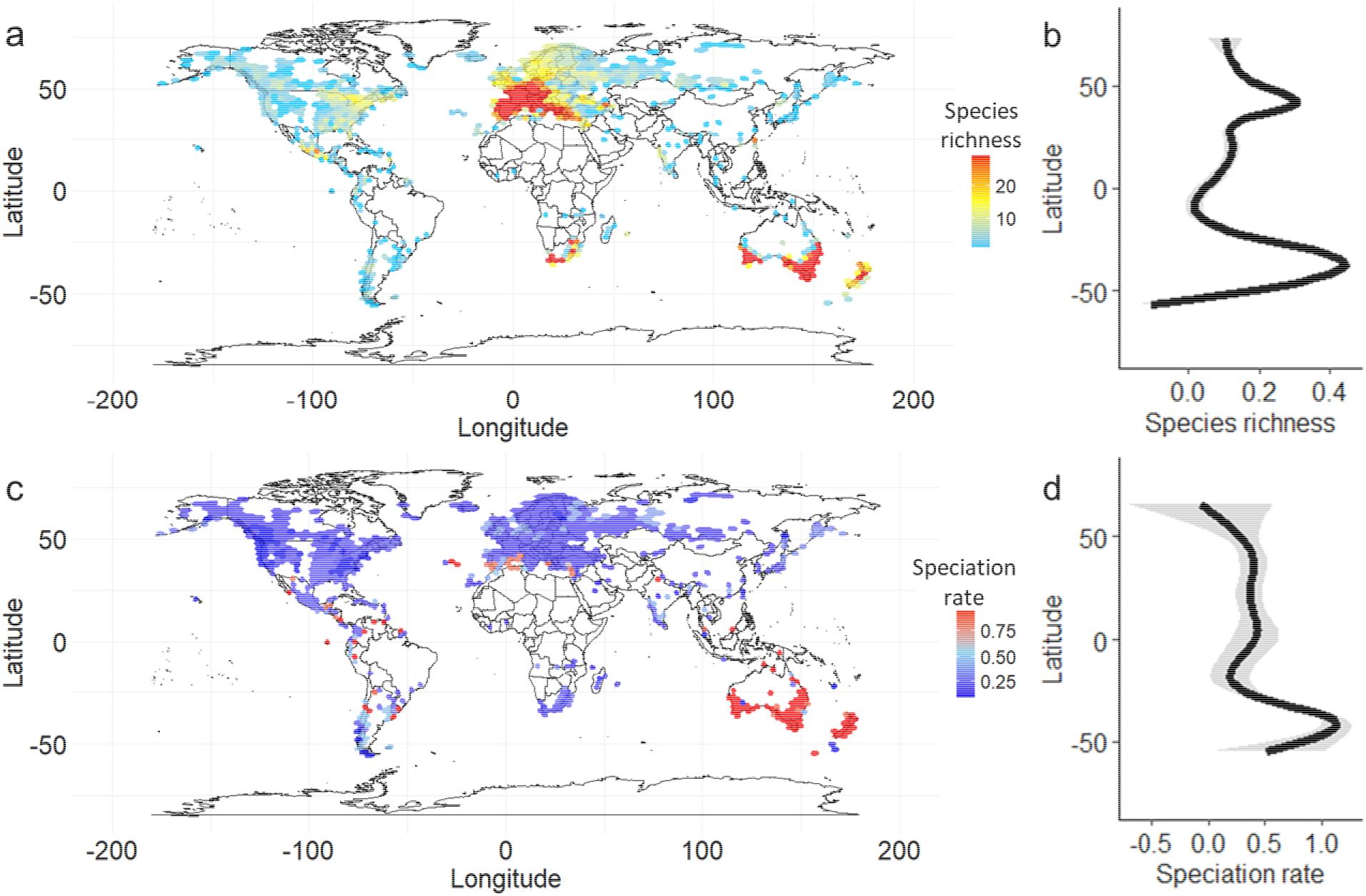
Global variation in orchidoid biodiversity. Species richness (**a**) and species richness by latitude (**b**) are calculated, respectively, from the number of orchidoid species represented in octagonal 200m^2^ grid cells spanning the earth and th**e** average number of species in occupied grid cells by latitude fit to a generalised additiv**e** model. Speciation rate (**c**) and speciation rate by latitude (**d**) were estimated by BAMM, respectively, within each grid cell and averaged across occupied grid cells by latitude f**it** to a generalised additive model.

Elevation has a similar apparent effect on speciation. Previous work indicates that orchid diversification can be promoted by cooler highland geolocations (41). We find apparent evidence for the opposite pattern in the orchidoids (SI Appendix, Fig. S5). However, tests of causality reveal no evidence of relationships between elevation and speciation rate (minimum elevation in species range: STRAPP p=0.42, Es-Sim p=0.29; maximum elevation in species range: STRAPP p=0.27, Es-Sim p=0.08).

### Worldwide cooling-driven speciation

Previous biogeographic research shows that orchids are best divided into seven geographical regions, defined as North America, Neotropics, Eurasia, Africa, Southeast Asia, Australia, Pacific (21, 45). We find that there are significant differences between mean speciation rates between some bioregions. In particular, Australia has significantly higher tip speciation rates as compared with all other bioregions (Africa: 0.1288; Australia: 0.2717; Eurasia: 0.1476; North America: 0.1346; Neotropics: 0.1623; Pacific: 0.2371; and Southeast Asia: 0.1653). However, as with latitude and elevation, there is no evidence for any causal relationship between speciation rate and bioregion (STRAPP, Kruskal-Wallis, p=0.085, SI Appendix, Fig. S6, Tab. S2-S4). Instead, we find consistent negative DCCA correlations between historic mean global temperature and speciation rate through time (Africa: −0.16; Australia: −0.57; Eurasia: −0.40; North America: −0.05; Neotropics: −0.34; Pacific: −0.42; and Southeast Asia: −0.48, p<0.0001 within each bioregion).

Our finding of cross-Earth influences of global cooling could be biased if taxa with high speciation rates are more likely to migrate between bioregions or if responses to climate change happen in taxonomically similar groups predominating each bioregion. By focusing on species endemic to each bioregion, we confirm that diversification happened independently and simultaneously across the earth. Though none are strictly monophyletic, the orchidoids species endemic within each bioregion are largely independent, as evidenced by phylogenetic clustering within each bioregion, except the Pacific, a species-poor sample (Africa D=-0.51, n=221; Australia D=-0.40, n=228; Eurasia D=-0.41, n=69; North America D=-0.51, n=29; Neotropics D=-0.57, n=109; Pacific D=0.28, n=33; Southeast Asia D=-0.03, n=47) (46). Importantly, we find that speciation rate increases with global cooling in 7 out of the 7 bioregions in independent fits of rate-through-temperature curves involving data spanning the last 10M years (Figure 5a). Shifts in speciation rate between bioregions are also positively temporally correlated (Figure 5b), providing strong support for independent, contemporaneous diversification driven by global cooling.

**Figure 5:**
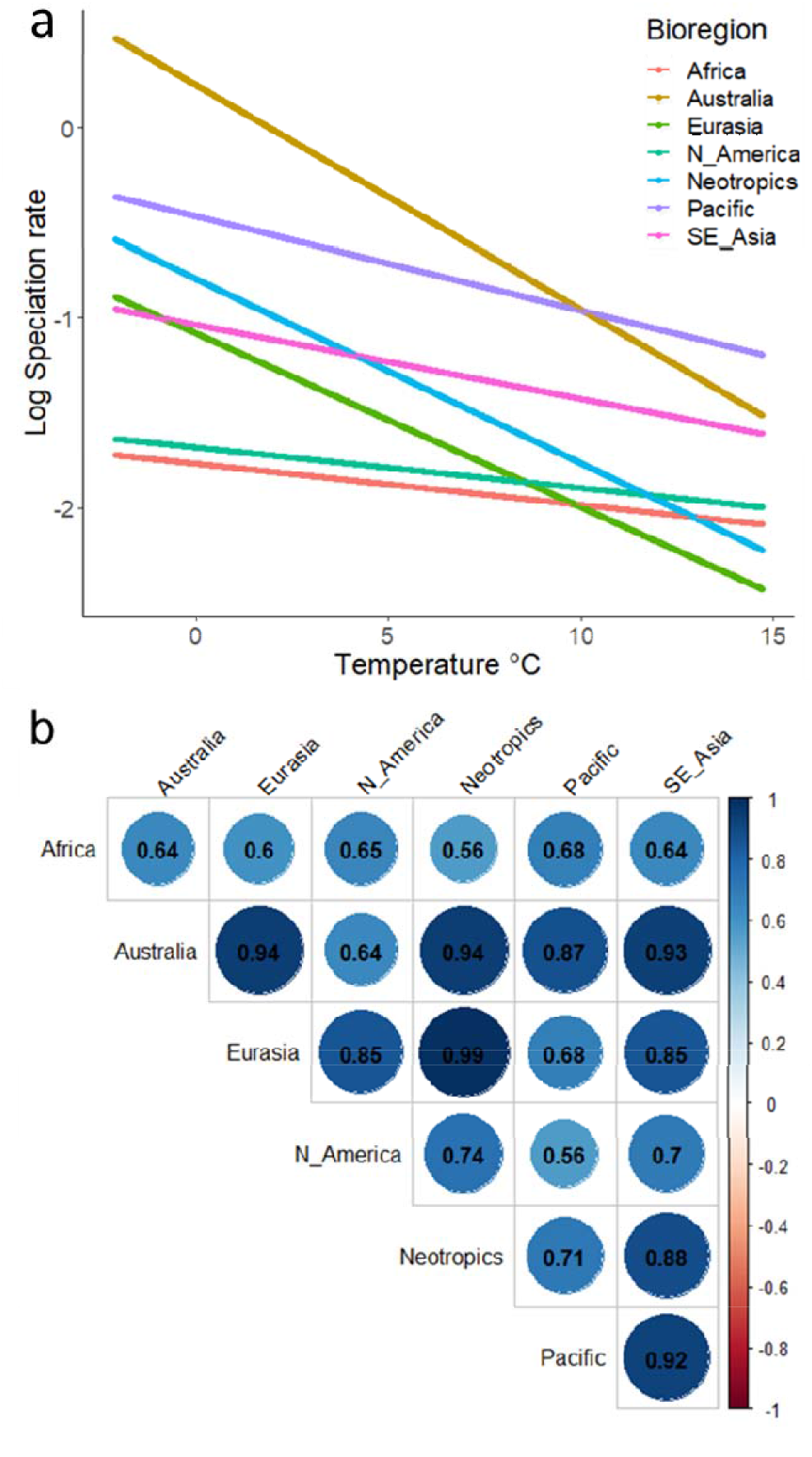
Geographic consistency in diversification driven by global cooling. Rate-through-temperature curves (**a**) of endemic lineages within each orchid bioregion were estimated by fitting exponential models to BAMM-derived speciation rate data covering the last 10M years. The correlation matrix of speciation rate through time curves (**b**) displays the degree to which diversification happened independently and simultaneously in endemic lineages across the Earth, from strong, +0.56 (light blue), to very strong, +0.99 (dark blue), correlations between endemic species in each bioregion.

### Diverse regional responses to other aspects of climate change

It is unlikely that geographic consistency in the influence of global cooling results from regression to the mean or insufficient power. Consistent with previous work (9–12), we find evidence that the type of climate change depends on the ecoregion in which it occurs (Table 1). Global atmospheric CO_2_ is positively correlated with speciation rate in two bioregions (Pacific and Southeast Asia); while global sea level is positively correlated with speciation rate in two different bioregions (Africa and North America; Table 1). Thus, though some aspects of mean global climate change are geographically varied, our study shows that global cooling drove speciation across the earth.

**Table 1:** Geographic variation in the consequences of mean global atmospheric CO_2_ and sea-level. DCCA correlations between speciation rate and climate indices are reported with significance indicated (* = p<0.05, ** = p<0.001).

| Bioregion | Atmospheric CO<sub>2</sub> | Sea-level |
| --- | --- | --- |
| Africa | -0.18 | 0.36** |
| Australia | 0.078 | 0.046 |
| Eurasia | -0.16 | 0.14 |
| Neotropics | -0.098 | 0.078 |
| North America | -0.27 | 0.57** |
| Pacific | 0.37** | 0.074 |
| Southeast Asia | 0.20* | 0.074 |

## Discussion

Though it is well recognised that conservation is important (47–51) and that research on the forces shaping biodiversity is critical to conservation (6, 52–58), it has been difficult to disentangle impacts of local climate from the historical consequences relating to time and global climate change. Previous models have predicted a general role for global climate change in biodiversity (7, 59), but the notion that climate change is responsible for how biodiversity is distributed across the earth has seemed far-fetched. Our study shows that global cooling is the major force underlying the biodiversity of terrestrial orchids. Our finding that biodiversity is geographically varied – despite being driven by climate change – demonstrates how ecological forces generating biodiversity can be concealed without considering evolutionary context, a finding with fundamental and applied implications.

Our findings provide an important counterexample to many theories of diversification. Though the orchidoids were not featured in the original debate on Darwin’s “abominable mystery” as they emerged well after the origin of angiosperms (1–5), our findings provide a clear case in which gradual diversification with time and singular origins of diversification are untenable. As speciation in the orchidoids is driven by global cooling, rather than warming, the findings run counter to classic explanations of rapid ecological niche filling (59) and the metabolic theory of biodiversity (60). We find no support for cradles of diversity, or highlands- or tropics-driven speciation (14, 17, 40, 41). And, our finding of consistent influences of historic temperature change runs counter to previous work describing variation in climate-driven diversification between closely related groups of taxa (8–10) or ecoregion (11,12).

Our findings beg the question of whether there are undiagnosed roles for climate change in other major radiations. Recent macroecology suggests that global cooling may have a general role in angiosperm speciation (57). The variation we found in 3 separate plant groups, shows that there is an urgent need for a meta-analysis of climatic effects on diversification in major plant groups, on par with that reported for four-legged animals (13). Still, for many highly specious plant taxa, such as cacti, *Euphorbia* and other succulent plants, it is difficult to test for effects of historic climate change because we are lacking needed paleoclimatic data sets such as aridity and other abiotic factors likely fundamental to their evolution. We predict that climate change will be associated with speciation in the largely epiphytic epidendroid orchids because, similar to their orchidoid sister clade, they exhibit recent explosive radiation, a consistent pollinia phenotype, and higher chromosome counts than earlier diverging orchid clades.

There is growing evidence of the influence of climate change on evolution. Recent work shows that the famous diversification of cichlid fish was driven by a temporally complex mixture of tectonic activity, climate change, biotic resource flow and interspecific hybridization (61,62). Rapid evolution in response to climate change is well-established in particular species of Darwin’s finches and Anolis lizards (63, 64). The persistence of ancient alleles in these groups (65, 66) is consistent with the presence of high genetic variability early in island colonisation; but, it is also in agreement with colonisation events followed by climate change-induced genetic variability. Though our study cannot differentiate between these models of evolution, it does highlight a need for detailed geolocation-specific records of historic climatic variation, as we found that the effects of global average CO_2_ and sea level are spatially varied consequences on speciation.

### Mechanisms of evolution driven by climate change

It is not clear how global cooling drives diversification. Milankovitch (orbital) cycles change the distribution of exposure to annual solar energy in predictable ways, but responses on Earth, including biotic changes, tectonic activity and climate change, can be varied (67). A role for tectonics is unlikely, as it is expected to have localised geographic impacts (67). The global expansion of C_4_ grasslands which peaked 4-8 Ma (68–71) could have contributed to more recent orchidoid speciation, but not the major inception of the burst in speciation.

Major oscillations in global temperature that occur during global cooling may have a critical role in accelerating orchidoid speciation, though evidence to date suggests global temperature oscillations are fairly recent (72). While the underlying mechanism is likely the same across the orchidoid tree of life, our finding that the Australian bioregion has the highest apparent speciation rate and exhibits the strongest evidence of relationships between global temperature and speciation rate could also be explained by climate change. As a consequence of the onset of the Antarctic Circumpolar Current (ACC) approximately 30 MYA (73), the Australian continent separated from Antarctica and moved north, resulting in dramatic cycles of cooling and drying experienced across Australia (74, 75). These repeated cooling cycles may have stimulated higher speciation rates as compared with other bioregions, the signature of which is preserved in the tip speciation rates seen today.

Global cooling could stimulate diversification by disrupting gene flow. Terrestrial orchids are sensitive to temperature variability (76, 77). Symbiotic mycorrhizal fungi of orchids provide cold-assisted improvements in germination (78), but also contribute to population subdivision (79). The tight coevolution between orchidiods and their pollinators first identified by Darwin (80, 81), also implies a rapid mechanism for generating barriers to gene flow. Indeed, pollination can be reduced through influences of climate change on plant phenology and physiological limitations of pollinating insects in colder temperatures (82).

Climatic cooling could drive rapid speciation by increasing genetic variability. Several studies show that biotic and abiotic factors, particularly temperature, can alter meiotic recombination in patterns which can accelerate adaptation (83, 84). Major changes in genome size and GC content are associated with orchidoid diversification (85, 86). Though we find that each of the major orchidoid subclades have more chromosomes than earlier evolving orchid taxa, without evidence of cooling-induced changes in chromosome number, the more reasonable hypothesis is that high chromosome numbers evolved early. Our finding of a well resolved orchidoid phylogeny with high speciation rates is consistent with speciation simultaneously occurring in reproductively isolated populations, perhaps as a consequence of climate-mediated stress (87), and, in rare cases, allotetraploidy events (88).

### Contemporary Implications

This research provides a clear case study of the long-term impacts of climate change on biodiversity. Preserving hot spots of diversity has been a central tenet of conservation, with substantial theoretical support (89). But, the logic of setting these conservation priorities hasn’t necessarily focused on the ecological processes that generate the biodiversity we are trying to conserve. If cross-earth speciation from climate change is common and unrelated to localised biodiversity, then conserving areas with low-species richness may be just as important for preserving evolutionary potential.

Resolving whether climate change stimulates adaptive radiation through stronger selection, population fragmentation or increased genetic variability has implications for conservation. There has been substantial interest in whether plants have sufficient standing genetic variation to respond to predicted gradual global warming (90). But, the history of earth is marked by catastrophic shifts in climate (91). And, climate change can impose strong breeding systems selection (ex. 92, 93). The relevant question for conservation could be whether plant species have sufficient capacities to generate novel genetic variation in response to climate-induced stress. Our finding that global cooling independently stimulated the formation of the thousands of orchidoid species within a short period (30 Ma) indicates that changes in climate can have predictable influences on speciation. Our study establishes terrestrial orchids as an excellent model system for understanding how climate-induced shifts in physiology fuel heritable variation, stimulate rapid adaptation and spur rapid speciation.

## Methods

### Supermatrix assembly

We mined Genbank for Orchidoideae sequences using the OneTwoTree pipeline (94), filtering intraspecific varieties, hybrids and open nomenclature. We corrected nomenclature against The Plant List (www.theplantlist.org), which reduces the impact of poor taxonomic assignment within Genbank. OneTwoTree clustered sequences into orthology groups, which we inspected and edited by reclustering partial sequences with full sequences. Although this could have resulted in unreliable alignments, OneTwoTree selected the longest sequence for every species, and we visually inspected resulting alignments. We downloaded outgroup sequences from Genbank, and aligned these with ingroup sequences using Mafft-add (95). After trimming unreliably aligned positions with trimAl-gappyout command (96), and concatenated alignments into a supermatrix using AMAS (97). Finally, we removed taxa with identical sequences, which are known to create short terminal branches that distort phylogenetic inference (98).

### Phylogenetic reconstruction

We produced a time-calibrated phylogeny in three steps. After an initial Maximum Likelihood (ML) search using RAxML V8 (99), we identified and removed taxa exhibiting rogue behaviour in the bootstrap (BS) replicates using RogueNaRok (100). This is an especially important procedure when using published sequences, reducing the impact of “chimeric taxa” created when sequences are misidentified and concatenated, which contain conflicting phylogenetic signal (101). We performed another ML search for the final molecular phylogeny used as input for divergence time analysis. ML searches were performed in the Cyberinfrastructure for Phylogenetic Research (102). Each ML search used 1,000 BS replicates to assess topological support, and applied an individual GTR model of nucleotide substitution to each locus partition. We enforced the monophyly of three clades in both searches: tribes Orchideae/Diseae, tribe Diurideae and tribe Cranichideae (see 20), in order to improve the likelihood calculation as in other large phylogenies (103). We calibrated the ML phylogeny against geological time using Penalized Likelihood with treePL (23). Orchids are poorly represented in the fossil record, with only one orchidoid fossil assigned with certainty, to subtribe Goodyerinae (104, 105). Instead we used robust secondary calibrations (19), constraining minimum and maximum ages for 15 major clades, according to the upper and lower bounds of 95% CIs. An initial treePL run primed parameters for the final analysis, in which multiple smoothing parameters were tested and cross validated.

### Speciation analysis

We reconstructed speciation history using the Bayesian Analysis of Macroevolutionary Mixtures (BAMM) framework (24), sampling four MCMC chains of 50 million generations every 5,000. We set priors with the R package BAMMtools (106) and used a conservative prior of one rate shift. Rate shifts were permitted in clades of more than five species, to improve convergence. Genus-level sample fractions accounted for incomplete sampling, which were derived from a checklist (20). We excluded the first 10% of generations as burn-in, and assessed convergence with the R package Coda (107), by confirming the effective sample size of each parameter was >200. We ignored reconstructions of extinction rate, which are known to be unreliable when modelled from phylogenies containing only extant taxa (25). We plotted the phylogeny against geological time with the R package strap (108).

We fit time- and temperature-dependent Maximum Likelihood models of diversification to the orchidoid phylogeny using the R package RPANDA (36), with code adapted from (107). Initially, we fitted 18 models starting from the simplest to models with increasing complexity (for each of time and temperature these included constant speciation with no extinction, linear speciation with no, constant, linear or exponential extinction, and exponential speciation with no, constant, linear or exponential extinction). We excluded from further analysis constant speciation models because of their poor explanatory power. We excluded linear speciation and extinction models because they are known to be problematic (108) and we find are biased against finding temperature-driven speciation (for example we find that 80% (66/82) of the taxa with best support (delta AIC<4) for temperature-driven speciation and extinction in a large study of tetrapods (13) exhibit non-linear model fits over relevant temperature ranges which prove a false impression of better fit to models of time than temperature). For direct comparisons of influences between temperature and time, we ultimately fitted 3 models (exponential speciation with no, constant, or exponential extinction) for both temperature and time to the orchidoid phylogeny by maximum likelihood. We calculated the corrected Akaike Information Criterion (AICc), the ΔAICc and the Akaike weight (AlCω) to assess likelihood support (see supplementary materials). We calculated the ΔAICc between specific models to test support for temperature and time, and plotted the best performing model against temperature and BAMM-estimated speciation rates. To compare the relative support for temperature vs. time across the three models, we calculated evidence ratios, estimated as (∑AICω temperature models / ∑AICω time models) for orchidoids in addition to published RPANDA parameters from 210 phylogenies (13) and estimated parameters for two published plant phylogenies (*Rhododendron* (37) and *Pinus* (38)). The distribution of the relative support for time-vs. temperature-driven diversification across the 213 phylogenies was displayed as violin plots of log-transformed evidence ratios to improve visualisation.

### Detrended cross-correlation analyses

We performed detrended cross-correlation analyses (DCCA) with the 9,001 post-burnin realisations of the speciation curve in R, using code from Davis et al (8). Briefly, we Tukey-smoothed paleoclimatic proxies, and interpolated values to the times recorded in the speciation rate curves. We calculated correlation coefficients between each of the 9,000 post burn-in speciation curves and paleoclimatic proxy, and plotted the distribution. A-1 indicates perfect negative correlation, 0 indicates no correlation, and +1 indicates perfect positive correlation (8). DCCA analyses were chosen over traditional correlation methods such as Pearson’s product moment correlation, because the time-series are autocorrelated over shorter timespans. A Wilcoxon rank sum test assessed significance of the distribution from the null hypothesis of zero (no correlation). Note, we did not include the monotypic tribe Codonorchideae in the analysis of cross-clade consistency. Paleoclimatic proxies of global mean temperature, atmospheric CO_2_ and sea-level were sourced from Zachos et al (29), Bergmann et al (109) and Miller et al (110), respectively.

### Chromosome counts

We downloaded available chromosome counts for all Orchidaceae from the Chromosome Count Database (33), using the median value where multiple counts were reported for a species.

### Biogeographical analyses

We downloaded georeferenced occurrences from GBIF (42) using rgbif (111), and cleaned coordinates with the R package CoordinateCleaner (112). We removed coordinates with uncertainty >10km, and those near capital cities, biodiversity institutions, those with equal longitude and latitude values, and within seas, resulting in >1.5 million cleaned records. To test for causal influences of spatial climatic variation, we used Structured Rate Permutations on Phylogenies (STRAPP) (43), Es-Sim (44) and phylogenetic signal calculated for binary traits with the D statistic (46) in the R package caper (113). In our analysis of tropical and nontropical speciation rates, two definitions of tropical and nontropical were used, a geographical definition and a temperature definition. The geographically-defined sample defined taxa as tropical if >50% of occurrences lie within -/+ 23.5 degrees of the equator. The temperature-defined sample assigned taxa as tropical if the year-round monthly temperature at >50% of occurrences exceeds 18°C (114,115), and to calculate this we retrieved data of global temperature (1970-2000) at 1km resolution, from WorldClim (116). These binary trait-dependent speciation analyses were performed with STRAPP, using 1,000 permutations and assessing significance with a Mann-Whitney test. In our analysis of speciation rates of major bioregions, we sorted taxa into seven bioregions defined by Givnish et al (21) (Africa, Australia, Europe, North America, the Pacific, and Southeast Asia), excluding taxa present in >1, and conducted another STRAPP test, using 1,000 permutations and assessed significance with a Kruskal-Wallis test. We used BAMMtools to estimate median speciation rate through time curves for each bioregion and tested temporal relationships between bioregions with Pearson correlations. In our analysis of elevation-dependent speciation, we acquired minimum and maximum elevation from the Internet Orchid Species Photo Encyclopaedia (http://orchidspecies.com/: accessed 31st July 2020), and assessed elevation-dependent speciation with STRAPP and Es-Sim, with both tests implementing 1,000 permutations. Es-Sim uses a Pearson test to assess significance, and we used a Spearman test in the STRAPP analysis. Elevation data from orchidspecies.com is likely to be partially qualitative, but elevation data in GBIF was poor. When performing DCCA analyses for regional speciation curves, we used all 9,001 post burn-in speciation curves when investigating the role of global cooling, but used mean speciation rate when correlating regions with each other, and with atmospheric CO_2_ and sea-level. We visualised variation in the relationship between speciation rate and temperature across bioregions in the most recent 10Ma, the timeframe with the most dramatic radiations. To do this, we fitted exponential models between historical speciation rate of bioregion-endemic species and paleotemperature (d18O) (29), and plotted the outcome.

## Acknowledgments

We thank the Younger laboratory (The Milner Centre for Evolution) for comments on earlier drafts. We thank the Darwin Correspondence Project and Riley-Anne Prydderch. We also thank Simon Pugh Jones, MBE, DSc of the Writhlington Orchid Project for advice and images. This study was supported by a Roger and Sue Whorrod Studentship (to J.B.T.), John Templeton Foundation Grant 61408 (to M.A.W.) and a Research England QR GCRF grant (to N.K.P.).

## Supplementary Information

https://datadryad.org/stash/share/t6xNcmKeDyLQDF1YFNEYfNaw0SWPsdIZMJnyx62IcYs

